# Testing retrogenesis and physiological explanations for tract-wise white matter aging: links to developmental order, fibre calibre, and vascularization

**DOI:** 10.1101/2024.01.20.576373

**Authors:** Tyler D. Robinson, Jordan A. Chad, Yutong L. Sun, Paul T. H. Chang, J. Jean Chen

## Abstract

To understand the consistently observed spatial distribution of white-matter (WM) aging, developmentally driven theories termed “retrogenesis” have gained traction, positing that the order of WM tract development predicts the order of declines. Regions that develop first are expected to deteriorate the last, i.e. “last-in-first-out”. Alternatively, regions which develop most rapidly may also decline most rapidly in aging, or “gains-predict-loss”. The validity of such theories remains uncertain, in part due to lack of clarity on the definition of developmental order. Importantly, our recent findings suggest that WM aging is also associated with physiological parameters such as perfusion, which may be linked to fibre metabolic need, which in turn varies with fibre size. Here we address the extent to which the degree of WM aging is determined by development trajectory (i.e. retrogenesis) and/or by physiological state. We obtained microstructural and perfusion measures using data from the Human Connectome Project in Aging (HCP-A), complemented by a meta-analysis involving maps of fibre calibre and macrovascular volume. Our results suggest that (1) while tracts that appear last or finish myelinating first in development display the slowest aging, the pattern of aging is not fully explained by retrogenesis; in fact, time courses of tract emergence and myelination give rise to opposite associations with WM decline; (2) tracts that appear earlier also have higher mean axon calibre and are also associated with lower degrees of WM microstructural aging; (3) such tracts also tend to exhibit relatively sustained CBF with a higher rate of lengthening of the arterial transit times (ATT), suggestive of collateral blood supply. These findings were also sex dependent in a tract-specific manner. Future work will investigate whether these are ultimately influenced by each tract’s metabolic demand and the role of macrovascular collateral flow.

## INTRODUCTION

There are well-documented spatial variations in the distribution of WM microstructural aging (Cox et al., 2016; de Groot et al., 2016; Michielse et al., 2010; Schilling et al., 2022; Slater et al., 2019), however, attempts to model the mechanism of these variations have produced only mixed success (Kiely et al., 2022; Slater et al., 2019; Yeatman et al., 2014). Developmentally driven models of brain aging pose an inverse temporal relationship between tract-wise white matter (WM) development and later declines in aging. These retrogenesis theories have gained considerable traction, positing that order or trajectory of tract-wise white matter (WM) development predicts later declines, sometimes simplified to suggest that degeneration occurs first in anterior, and last in posterior regions, i.e. “last-in-first-out (LIFO)” (Bender et al., 2016; Hoagey et al., 2019; Raz, 2000). However, the validity of LIFO theories remains uncertain in microstructural research, in part due to lack of clarity in defining first or last “in” in the context of numerous measures of neural development. In fact, irrespective of which tract is first to appear prenatally, many regions of the brain continue to mature into adulthood (Lebel & Beaulieu, 2011; Yeatman et al., 2014). Another theory, termed “gain-predicts-loss” (Slater et al., 2019), posits that the rate of development predicts the rate of later declines, in which those regions which develop most rapidly will also decline most rapidly in aging. However, there are again several metrics by which the rate of development can be asserted (Grotheer et al., 2022; H. Huang et al., 2006; Kiely et al., 2022; Yu et al., 2020).

One of the earliest markers available for defining a tract as “in” in the context of a developmentally ordered last-in-first-out model is the point at which a tract becomes recognizable in prenatal development (H. Huang et al., 2009; Y. Ouyang et al., 2021). Prior to the emergence of a tract as a distinct structure in the prenatal brain, meaningfully differentiating regions for comparison to adult tracts becomes conceptually challenging. As one of the earliest possible ways to define a tract’s arrival, prenatal emergence thus represents a valuable starting point for examining retrogenesis predictions. The degree to which this stage of development can be considered a foil for WM degeneration in aging remains unclear, as the earliest stages of tract development may have less in common with age-related degeneration than later and ongoing developmental processes (Brody et al., 1987; Grotheer et al., 2022). While WM microstructural development continues into adulthood (Arshad et al., 2016; Slater et al., 2019; Stricker et al., 2009), the degree of myelination at birth as defined by R1 values may represent a proxy measure for developmental state in the context of retrogenesis under the assumption that those tracts showing highest myelin density at birth are those which begin myelinating earliest (Grotheer et al., 2022).

While last-in-first-out models of retrogenesis can be addressed via early WM developmental states, addressing gain-predicts-loss hypotheses into adulthood requires incorporating ongoing rates of WM maturation. Peak microstructural integrity across the lifespan has been posited as a measure of the full maturation of a region (Slater et al., 2019; Yu et al., 2020). However, tract-wise ages of peak mean diffusivity (MD) do not mirror the order in which tracts reach peak R1 values indicating maturation of myelin packing (Yeatman et al., 2014), necessitating separate predictive models representing different developmental processes. Notably, myelination state at birth and changes in myelination into adulthood also demonstrate differing orders by tract (Grotheer et al., 2022; Slater et al., 2019).

In addition to early developmental orders and rates of tract maturation, physiological determinants such as axon calibre have been considered a measure of WM maturation, as increasing axon calibre is a notable component of early WM fiber development and associated with brain function (Genc et al., 2018; S. Y. Huang et al., 2020). Our recent findings suggest that WM degeneration may also vary by both axonal size and perfusion (Robinson et al., 2023). As a result, incorporating physiological differences between tracts can be considered both a component of, and a complement to, developmental retrogenesis models. As thicker fibres are also thought to be more heavily myelinated and more efficient conductors (Mancini et al., 2021; Perge et al., 2012), the relationship between the aging trajectory, fibre calibre and blood supply could help shed light on the broader mechanisms governing this trajectory. For the purposes of this study, we have targeted mean axonal calibre and macrovascular density as well as perfusion as relevant physiological determinants for tract-wise differences in WM aging.

The degree to which WM degeneration in aging is determined by development trajectory or physiological factors is central to understanding the mechanisms of WM degeneration. Using diffusion MRI (dMRI) and perfusion data drawn from the Human Connectome Project in Aging (HCP-A) data set, this study aims to test tract-wise retrogenesis predictions in WM integrity across six models of WM development and maturation drawn from existing literature to determine which, if any, best represent degrees and rates of decline in the aging WM. Additionally, we aim to demonstrate that the trajectory of WM decline may hinge on not only development, but also on physiological attributes that govern WM function, such as fibre calibre, vascularization and perfusion.

## METHODS

### Data sets

#### White matter microstructure and perfusion

535 healthy adult subjects (235 male and 300 female, aged 36-100, approx. 48% postmenopausal) were drawn from the HCP-A dataset (OMB Control# 0925-0667) (Bookheimer et al., 2019; Harms et al., 2018). All subjects were assessed to be in good health and without pathological cognitive impairment (i.e. stroke, clinical dementia). Manual quality control identified thirty-two subjects for exclusion from the original sample due to MRI artifacts that produced notable errors when generating WM masks. Three additional subjects were excluded as age outliers due to being the only subjects of 100 years of age, bringing the final sample size to 500 subjects (206 male and 294 female, aged 36-89) (**Table 1**).

**Table 1:** Sample Demographics. Demographic statistics for all subjects, female subjects, and male subjects. All subjects were in good health and without pathological cognitive impairment (i.e. stroke, clinical dementia)

|  |  |  |  |  |  |  |  |
| --- | --- | --- | --- | --- | --- | --- | --- |
|  | Count | Mean Age (years) | Median Age (Years) | Age Range (Years) | Age Stdev (Years) | Hypertension Count | Mean Education (years) |
| **All** | **500** | **57.88** | **56** | **36-89** | **14.3** | **134** | **17.41** |
| Female | 294 | 57.27 | 54 | 36-89 | 14.4 | 63 | 17.29 |
| Male | 206 | 58.76 | 58 | 36-89 | 14.2 | 71 | 17.58 |

The study accessed T1-weighted structural MRI, diffusion-weighted MRI (dMRI), with a 1.5mm^3^ voxel resolution, MB=4, with 93 directions at b=1500 and 3000 s/mm^2^, and 6 b=0), and multi-delay multi-band echo planar imaging (MB-EPI) pseudo-continuous arterial spin labeling (pCASL) with a 2.5mm^3^ voxel resolution, MB=6, with TIs = 0.7, 1.2, 1.7, 2.2, and 2.7s. Data were collected using four matched Siemens Prisma 3T MRI scanners. Further imaging parameters for structural, diffusion, and perfusion imaging can be found in (Harms et al., 2018).

##### Fibre calibre and vascular density

Tract-wise axon calibre measures used in physiological determinant models and tractwise correlations in this study were derived from data analyzed by Gast et al. using axonal spectrum imaging (AxSI) from a 324 subject healthy adult sample (22-37 years of age, mean age 28.9, 180 female, 144 male) drawn from the HCP young adult dataset. Subjects were scanned on a 3T Magnetom Siemens Skyra scanner with a 128-channel RF coil and customized SC72 gradient system reaching 80mT/m using multi-shell diffusion-weighted imaging with Δ/δ = 43.1/10.6 [ms] and b-shells of 1000, 2000 & 3000 [s/mm2], as well as an MPRAGE sequence with TR/TE = 2400/2.14 [ms] (Gast et al., 2023). Axon calibre maps provided by Gast et al., which are available publicly at https://github.com/HilaGast/AxSI, were registered to tract masks generated by this study to produce mean tract-wish axon calibre values across subjects. Tract-wise macrovascular density values were calculated using data provided by Bernier et al of forty-two neurologically healthy subjects (22 years of age (20-31)) (Bernier et al., 2018). Images were acquired using whole-brain multi-band ToF angiography (200 × 200 × 120 FOV, TR/TE 23/3.6 ms, voxel size of 0.625 × 0.625 × 1.3 mm) venous inflow suppression, and a high-resolution multi-echo SWI sequence (230 × 230 × 160 FOV, TR 28 ms, TE 6.9/12.6/18.3/24.0 ms, voxel size of 0.6 × 0.6 × 1.2 mm). In-house preprocessing of ToF, SWI, and VED can be found in Bernier et al,’s respective methods section. Combined SWI and VED values were used to determine density values as the likelihood of vessels within a given voxel.

### Image analysis

dMRI data from the HCP-A data set were corrected for eddy-current and susceptibility-related distortions via EDDY (Andersson & Sotiropoulos, 2016), and eighteen major WM tracts were reconstructed from the dMRI data using FreeSurfer TRACULA (v 7.2.0) (Maffei et al., 2021; Yendiki et al., 2011), then combined bilaterally to produce ten tracts of interest (8 combined across hemispheres and 2 comissural) for analysis in local space using FSL’s fslmaths function. Resulting tracts included the major forceps (Fmajor), minor forceps (Fminor), anterior thalamic radiation (ATR), cingulum angular bundle (CAB), cingulate gyrus (CCG), corticospinal tract (CST), inferior longitudinal fasciculus (ILF), superior longitudinal fasciculus parietal (SLFP), superior longitudinal fasciculus temporal (SLFT), and the uncinate fasciculus (UNC) (**Figure 1**). This choice of tract segmentations was made to minimize spatial overlap between tracts.

**Figure 1:**
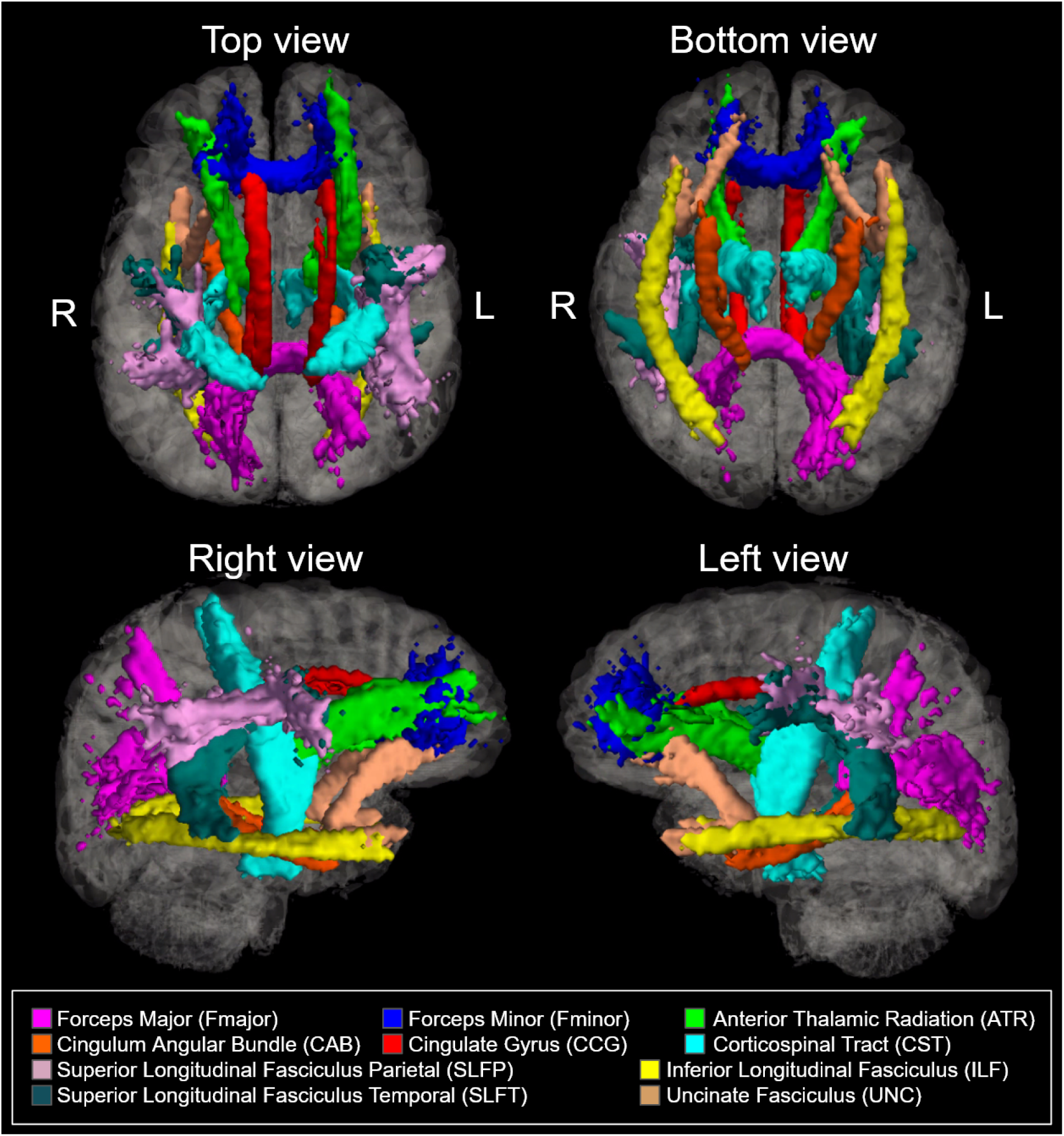
Ten bilateral tracts of interest reconstructed using Freesurfer’s TRACULA package. This collection of tracts of interest was selected to limit overlap between tracts of interest.

#### Microstructure

Microstructural integrity in WM tracts was quantified using fractional anisotropy (FA), mean diffusivity (MD), axial diffusivity (AD), and radial diffusivity (AD) measures. FA, MD, AD, and RD maps were derived via Dipy’s DKI script to achieve kurtosis-corrected DTI metric fitting due to the high b-value used in the HCP-A acquisitions. Per-subject tract masks generated by TRACULA were applied to local dMRI images in fslmaths to generate tract means for FA, MD, AD, and RD respectively. For each metric, baseline normalized (percent of baseline) values were calculated as individual raw tract values divided by the mean raw tract value of the youngest 10% of subjects to assess percent differences from the youngest baseline, resulting in “FA_perc_”, “MD_perc_”, “AD_perc_”, “RD_perc_” values used in subsequent analyses.

#### Perfusion

Cerebral blood flow (CBF) was quantified using the general kinetic model that accounts for arterial-transit delay (ATT) (FSL oxford_ASL) (Chappell et al., 2009, 2011) to produce per-subject maps of both CBF and ATT in local space. As with microstructural measures, TRACULA-generated tract masks were applied to the local maps using fslmaths to produce tract means for raw CBF and ATT. Both were corrected for partial-volume effects with the grey matter. Baseline normalized perfusion values were again calculated as individual raw tract values divided by the mean raw tract value of the youngest 10% of subjects (i.e. the baseline group) to produce “CBF_perc_” and “ATT_perc_” measures for use in subsequent analyses.

### Statistical analysis

Tract orders tested in each analysis were determined using six ordered models drawn from existing literature, as summarized in **Table 2**:

**Table 2:** Model definitions and corresponding tract order directionalities.

|  |  |  |
| --- | --- | --- |
| **Model** | **Tract ranking** |  |
|  | **Low** | **High** |
| **LAFO**<br>(last appearing first out) | First appearing prenatally | Last appearing prenatally |
| **LMFO**<br>(last myelinating first out) | First myelinating prenatally | Last myelinating prenatally |
| **SPFO**<br>(slowest peaking first out) | Lowest rate of MD change from adolescence to adult peak | Highest rate of MD change from adolescence to adult peak |
| **SMFO**<br>(slowest myelinating first out) | Lowest rate of R1 change from adolescence to adult peak | Highest rate of R1 change from adolescence to adult peak |
| **ADD**<br>(axon calibre (AxD) determined) | Lowest axon calibre | Highest axon calibre |
| **VDD**<br>(vascular density (VascD) determined) | Lowest macrovascular density | Highest macrovascular density |

#### Last-in-first-out models

(1) “Last-appearing-first-out” (LAFO): ordered by the time of prenatal emergence (Y. Ouyang et al., 2021). The ordinal variable comprises the order of earliest to latest prenatal emergence, including for the ATR, CCG, CST, Fminor, UNC, CAB, ILF, Fmajor, and SLF (combined SLFP/SLFT) in order of earliest to latest prenatal emergence.
(2) “Last-myelinated-first-out” (LMFO): ordered by R1 (“myelination”) at birth (Grotheer et al., 2022). The order variable comprises the order of lowest to highest degree of myelination at birth, including for the CST, ATR, SLF (combined SLFP/SLFT), UNC, ILF, Fmajor, CCG, and Fminor in order of lowest to highest degree of myelination at birth.

#### Gain-predicts-loss models

(3) “Slowest-peak-first-out” (SPFO): continuous variable that reflects the rate of MD change from age 15 to adult peak using quantitative tractwise mean rate of MD change values (Slater et al., 2019). Values are given as rates of MD change, including for the Fmajor, Fminor, SLF (combined SLFP/SLFT), CST, CCG, ATR, ILF, and UNC in order or lowest to highest rate of MD change before maturation.
(4) “Slowest-myelinated-first-out (SMFO): continuous variable that reflects the rate of R1 change from age 15 to adult peak using quantitative tractwise mean rate of R1 change values (Slater et al., 2019). Values are given as the rate of myelination, including for the SLF (combined SLFP/SLFT), Fmajor, Fminor, CST, CCG, ILF, ATR, and UNC in order of lowest to highest rate of myelination.

#### Physiological determinant models

(5) “Axon-calibre-determined” (ADD): continuous variable that reflects the axon calibre using quantitative tractwise mean calibre values (Gast et al., 2023). Values are given as the mean axon calibre (AxD), including for the CAB, UNC, ILF, ATR, Fminor, Fmajor, CCG, SLFT, SLFP, and CST in order of lowest to highest mean axon calibre.
(6) “Vascular-density-determined” (VDD): continuous variable that reflects macrovascular density (VasD) obtained from quantitative tractwise mean vascular density values (Bernier et al., 2018). Values are given as the mean macrovascular density, including for the CCG, SLFP, SLFT, ATR, UNC, ILF, Fminor, Fmajor, CST and CAB in order of lowest to highest mean macrovascular density.

Relationships between tract order, age, and tract-wise measures of mean WM microstructure and perfusion percent differences from baseline were assessed using multivariate regression in R (version 4.1.1). Tract-wise age effects were assessed using linear regression of age onto FA_perc_, MD_perc_, CBF_perc_, and ATT_perc_. Rosner’s tests were performed to identify and exclude volume outliers per tract in single tract analyses. The resulting subject counts for these analyses were: Fmajor (N=493), Fminor (N=493), ATR (N=497), CAB (N=490), CCG (N=495), CST (N=491), ILF (N=497), SLFP (N=497), SLFT (N=495), and UNC (N=495).

Retrogenesis predictions were addressed via tract-wise linear regressions were performed in terms of both the tract “state” and “rate” values. That is, under the tract ordering or respective values in each of these models (**Table 3**), we assessed the association between the tract’s rank in the given model’s and the mean tract parameters (“state-based”) as well as the association with the regression coefficient against age (“rate-based”), with sex as a covariate of no interest for computing the regression coefficient against tract ranking. Tract parameters include both microstructural (FA_perc_, MD_perc_) and perfusion (CBF_perc_, ATT_perc_) measures. Since they are normalized by baseline values, the averaged parameters provide insight into whether the tract begins its decline early or late.

**Table 3:** Tract values by model.

| Table 3: Tract values by model |  |  |  |  |  |  |
| --- | --- | --- | --- | --- | --- | --- |
| Tract | Model |  |  |  |  |  |
|  | Ordinal |  | Continuous |  |  |  |
| | LAFO | LMFO | SPFO(%) | SMFO (%) | ADD ($\mu\text{m}$) | VDD (%) |
| **Fmajor** | 8 | 6 | -1.92 | 1.80 | 4.70 | 1.79 |
| **Fminor** | 4 | 8 | -2.45 | 2.30 | 4.25 | 1.26 |
| **ATR** | 1 | 2 | -3.54 | 4.74 | 4.16 | 0.53 |
| **CAB** | 7 | n/a | n/a | n/a | 2.00 | 2.90 |
| **CCG** | 2 | 7 | -3.24 | 3.53 | 4.99 | 0.51 |
| **CST** | 3 | 1 | -2.75 | 2.88 | 5.90 | 2.61 |
| **ILF** | 7 | 5 | -3.56 | 4.05 | 3.89 | 0.88 |
| **SLFP** | 9 | 3 | -2.49 | 1.71 | 5.90 | 0.51 |
| **SLFT** | 9 | 3 | -2.49 | 1.71 | 5.85 | 0.53 |
| **UNC** | 5 | 4 | -4.51 | 6.54 | 2.65 | 0.85 |

#### Baseline correlations

Baseline associations, i.e. associations from within the youngest 10% of the sample, were examined between all possible pairs of variables both within and between whole-WM mean microstructural (FA_perc_, MD_perc_, AD_perc_, RD_perc_), perfusion (CBF_perc_, ATT_perc_), and physiological (axon calibre (AxD), macrovascular density (VasD)) measures. Pearson’s correlation coefficients were calculated for each variable combination in Matlab. Subject-wise correlations were conducted between all microstructure and perfusion values, while additional tract-wise correlations were conducted in both microstructure-perfusions comparisons and in correlations using axon calibre or macrovascular density values. Tract-wise correlations were used in lieu of subject-wise correlations due to the tract-wise nature of available physiological measures and mean tract-wise microstructural and perfusion measures were substituted for the baseline corrected per-subject values used in previous comparisons.

#### Effects of sex

Sex differences in the effects of tract order on mean microstructural (FA_perc_, MD_perc_, AD_perc_, RD_perc_) and perfusion (CBF_perc_, ATT_perc_) measures were conducted per model using multivariate regression of tract order, sex, and the interaction effect of sex by model-specific tract developmental order (LAFO, LMFO), maturation measure (SPFO, SMFO), or physiological measure (ADD, VDD) onto each respective microstructural/perfusion measure. Similarly, sex differences in the effects of developmental order, maturation measure, or physiological measure on rates of microstructural (FA_perc_, MD_perc_, AD_perc_, RD_perc_) and perfusion (CBF_perc_, ATT_perc_) declines per model with age were assessed via multivariate regression of tract order, age, sex, the interaction of development order, maturation measure, or physiological measure by sex, and the three-way interaction of tract order, age, and sex onto each respective microstructural/perfusion measure.

## RESULTS

Age-related declines were observed in MD_perc_ for all ten tracts, and in FA_perc_ for seven of ten tracts (Fmajor, Fminor, ATR, CAB, CCG, ILF, and SLFP; not CST, SLFT or UNC). Eight of ten tracts (Fmajor, ATR, CAB, CST, ILF, SLFP, SLFT, UNC; not CCG or Fminor) demonstrated significant age-related increases in AD_perc_, while significant declines in AD_perc_ were identified in the CCG alone. All ten tracts showed significant age-related increases in RD_perc_. Age-related declines in CBF_perc_ were identified in eight of ten tracts (Fmajor, ATR, CAB, CST, ILF, SLFP, SLFT, and UNC; not CCG or Fminor), along with age-related increases in ATT_perc_ in seven of ten tracts (Fminor, ATR, CCG, CST, SLFP, SLFT, and UNC; not CAB, Fmajor or ILF) (**Figure 2**).

**Figure 2:**
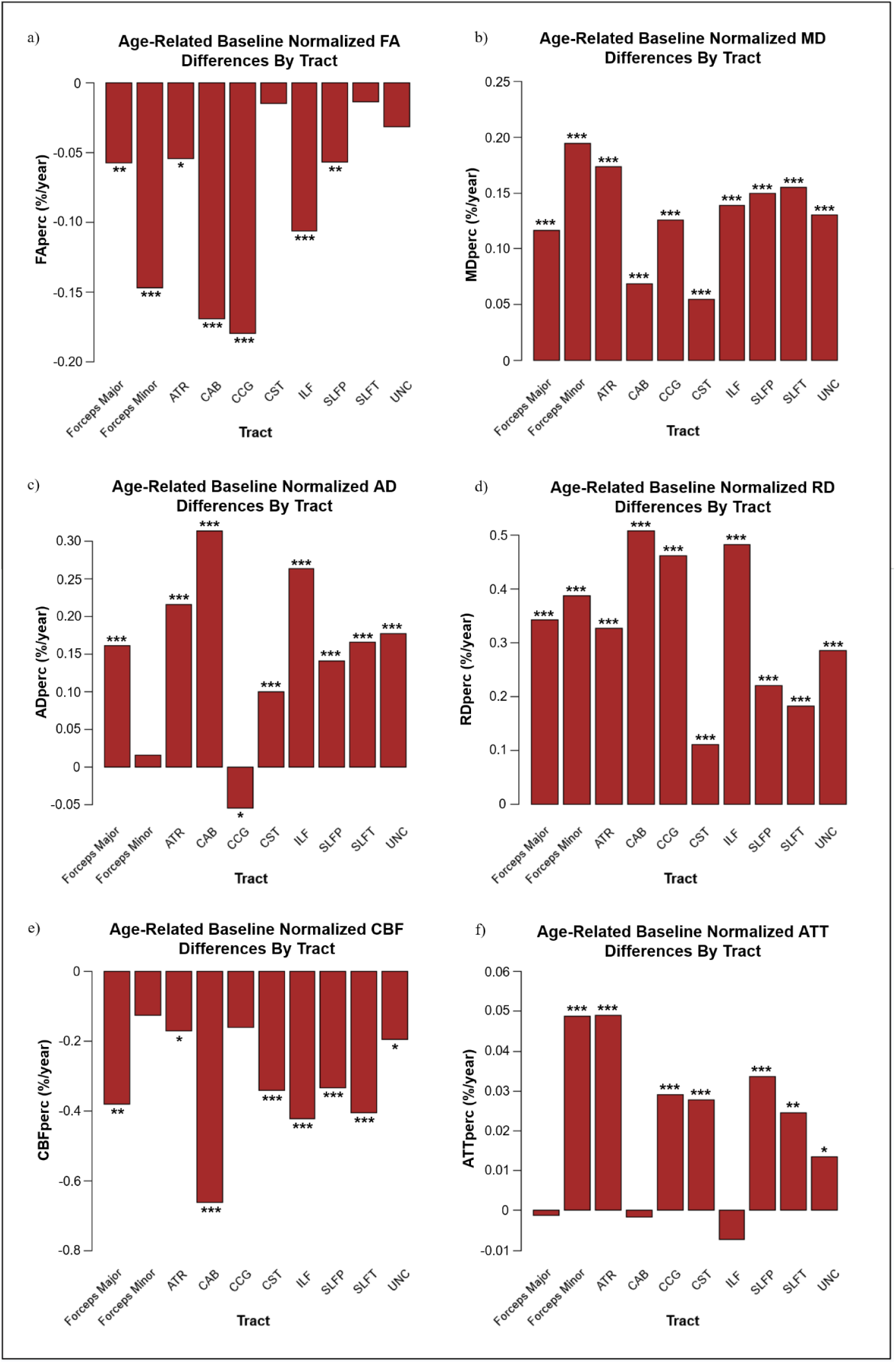
Tractwise age effects for FA**_perc_** (a), MD_perc_ (b), AD_prec_ (c), RD_prec_ (d), CBF_prec_ (e), and ATT_prec_ (f) respectively. Significant effects by sex ana age are denoted with asterisks (***: p<.001. **: p<.01, and *: p<.05).

### Order of state of microstructure declines

#### Last-in-first-out models

Both last-in-first-out models (LAFO and LMFO) demonstrated significant associations with at least one measure of microstructural integrity, with the LAFO model defined as greater declines in earlier appearing tracts and the LMFO model defined as greater declines in those which myelinate latest. In the LAFO model, tract development order was positively associated with mean FA_perc_ and AD_perc_, while no significant association with MD_perc_ or RD_perc_ was observed. In LMFO, FA_perc_ was found to be inversely related to R1 order, while MD_perc_ was positively associated with tract order. AD_perc_ and RD_perc_ also revealed significant associations with LMFO tract order, such that AD_perc_ was negatively associated with myelination order while RD_perc_ was positively associated (**Figure 3a,d,g**).

**Figure 3:**
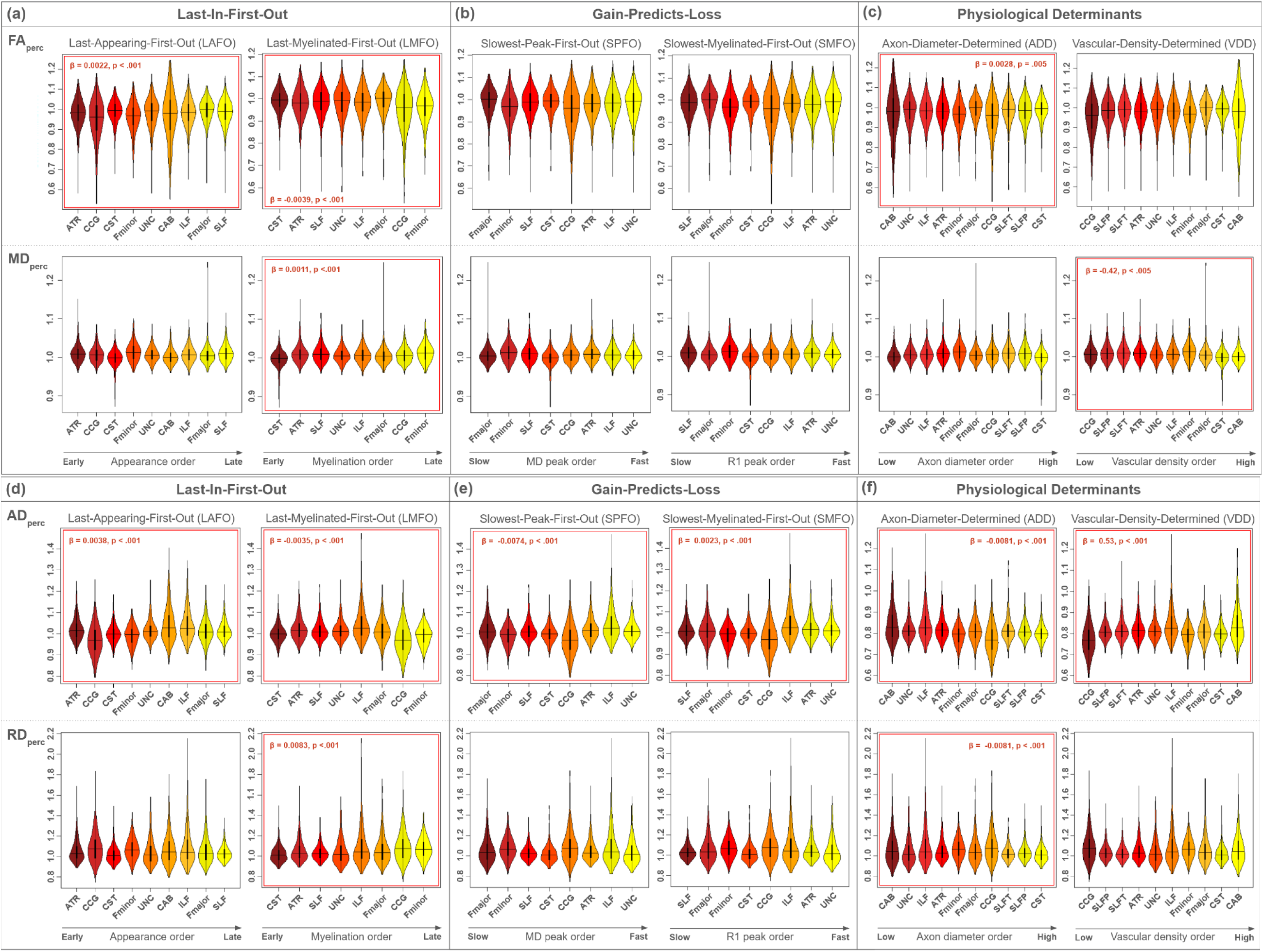
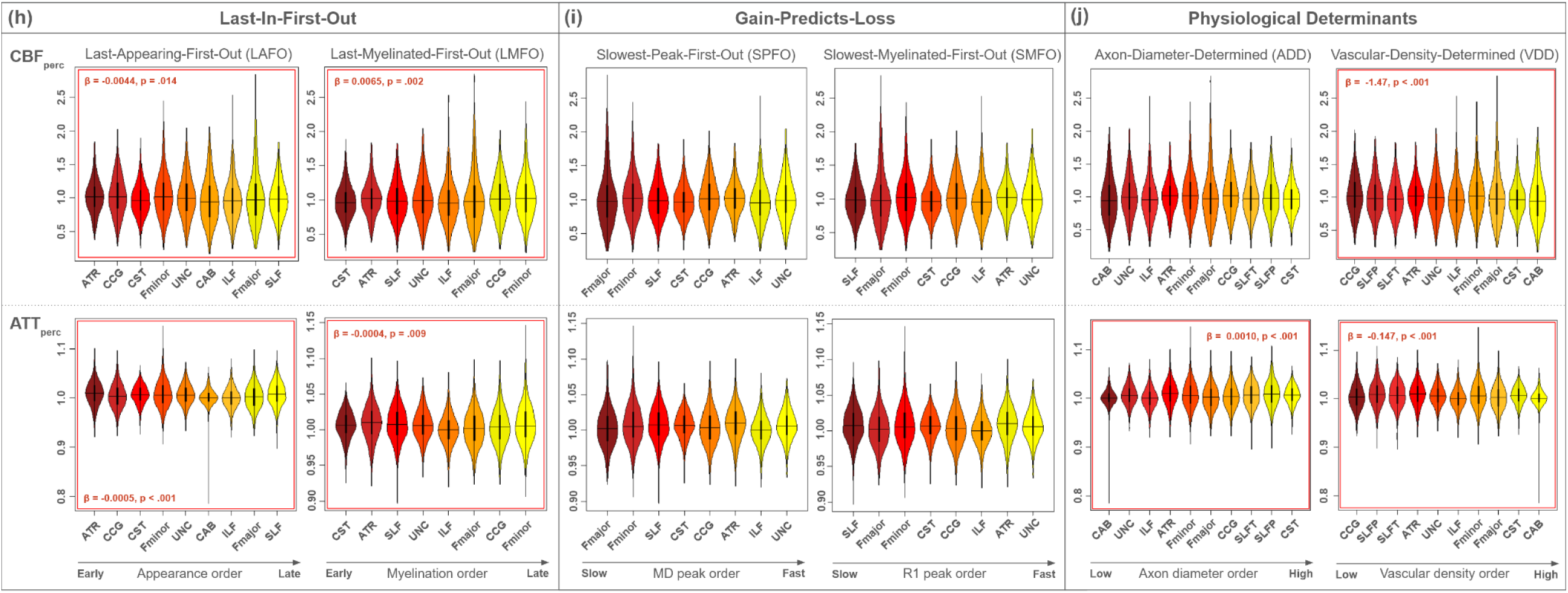
Tract-wise distributions for FA_prec_,MD_prec_, AD_prec_, RD_prec_, CBF_prec_, and ATT_prec_ under each Last-In-First-Out (a). Gain-Predicts-Loss (b), and Physiological Determinant (c) model. Significant linear relationships between tract order and microstructural/perfusion measure are denoted with a red outline.

#### Gain-predicts-loss models

Regarding gains-predicts-loss predictions, AD_perc_ was the only related variable. AD_perc_ demonstrated the only significant associations in mean microstructural measures, with AD_perc_ associating positively with rate of myelination under the SMFO model as faster myelinating tracts demonstrated the highest microstructural values, and negatively under the SPFO model as faster MD maturation was associated with lower AD_perc_ values (**Figure 3b,e,h**).

#### Physiological determinant models

Under the ADD model, a positive association was identified between axon calibre and FA_perc_ while a negative association was identified with AD_perc_ and RD_perc_. As for VDD, VasD was negatively associated with MD_perc_ but positively associated with AD_perc_, and nothing else.(**Figure 3c,f,i**).

### Order of rates of microstructural decline

In **Figure 4**, for each of the 6 models of tract ordering, we summarize the linear fits of each tract microstructural parameter against age in the colour maps. Bright colours represent higher values in a given microstructural parameter, and the regression coefficients for the age associations are plotted against tract rankings for each model (**Figure 5**). The result allows us to assess the association between tract ranking under each model and the rate of age-related tract decline. Significant regressions are indicated by red lines at the p<0.05 level

**Figure 4:**
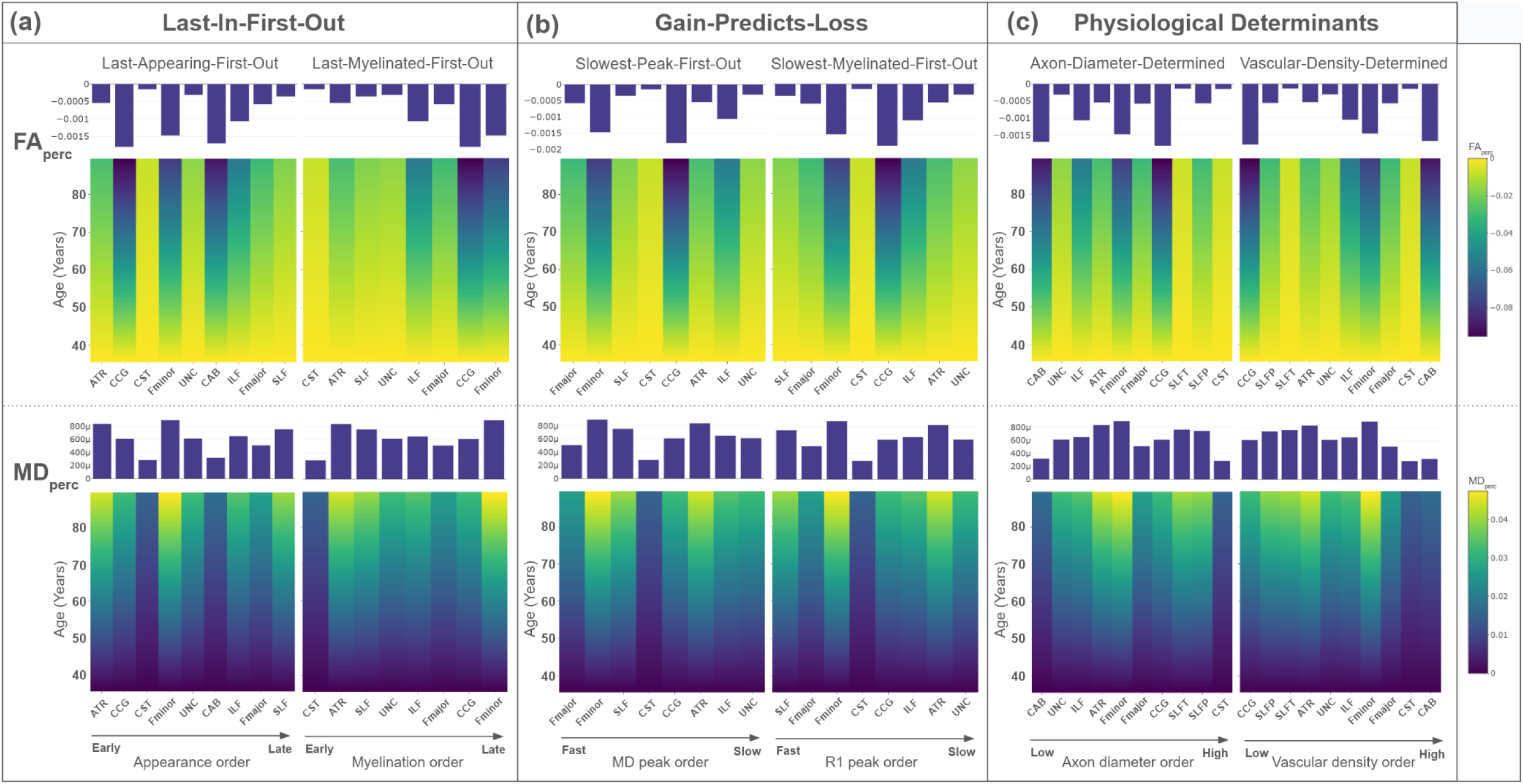
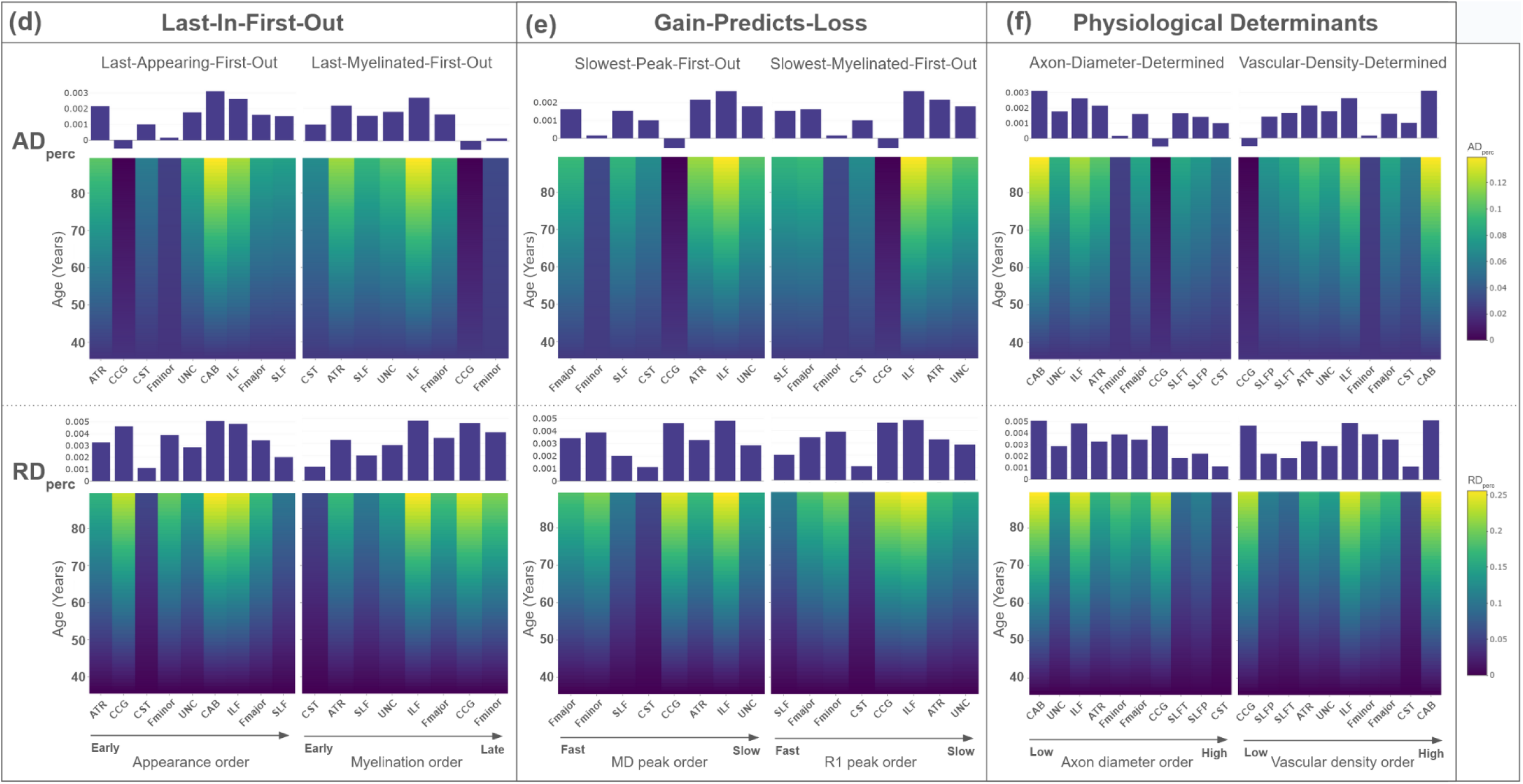
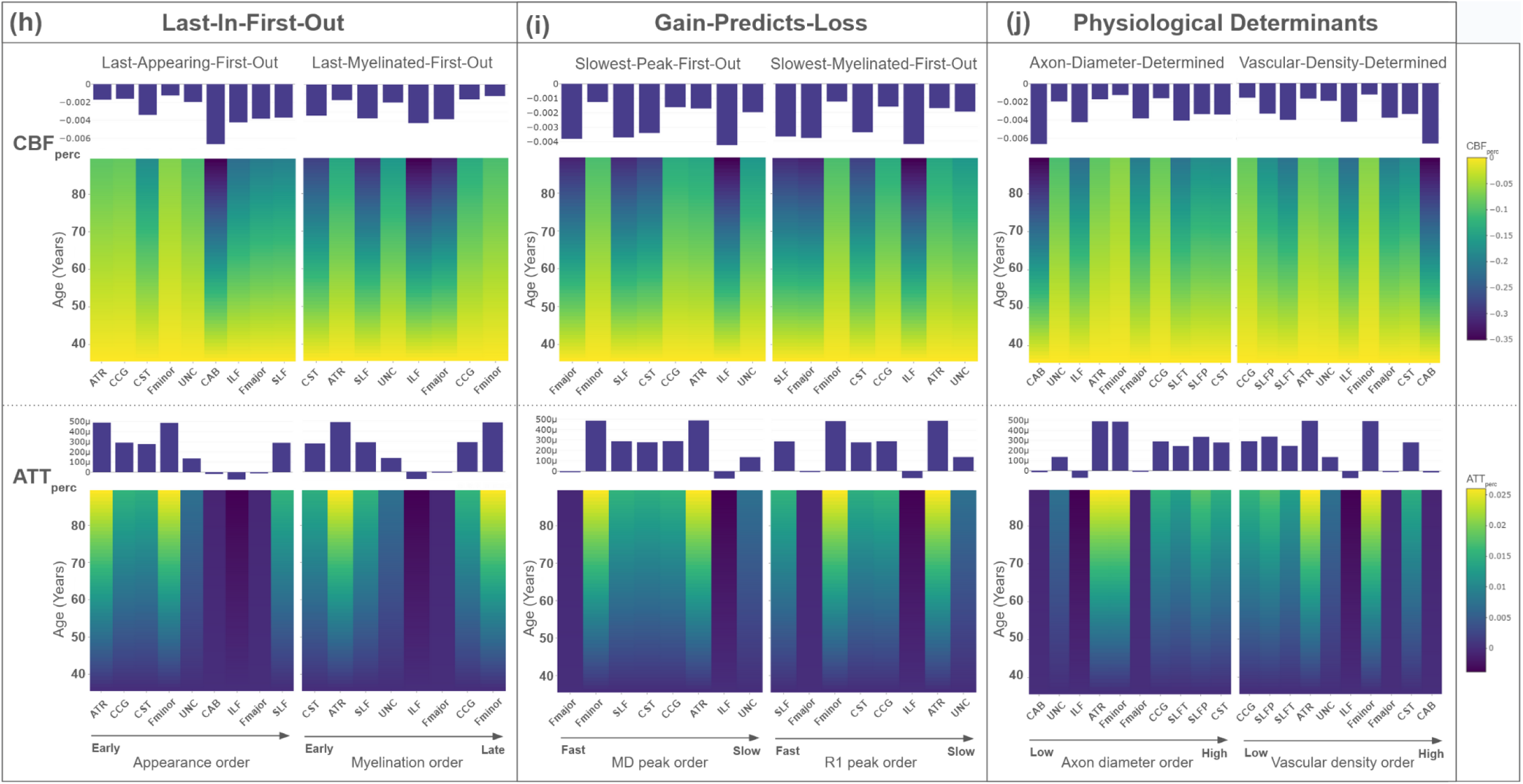
Tractwise age-effects bi FA_prec_, MD_prec_, AD_prec_, RD_prec_, CBF_prec_, and A TT_prec_ under each Last-hi-First-Out (a.d.h). Gain-Predicts-Loss (b.e.i) and Physiological Determinant (cjj) model, hi each plot, the colours in each column indicates the values of the variable of interest modeled as a linear function of age, and bright colours represent higher values. Rates of age-related change per tract are plotted in corresponding line graphs.

**Figure 5:**
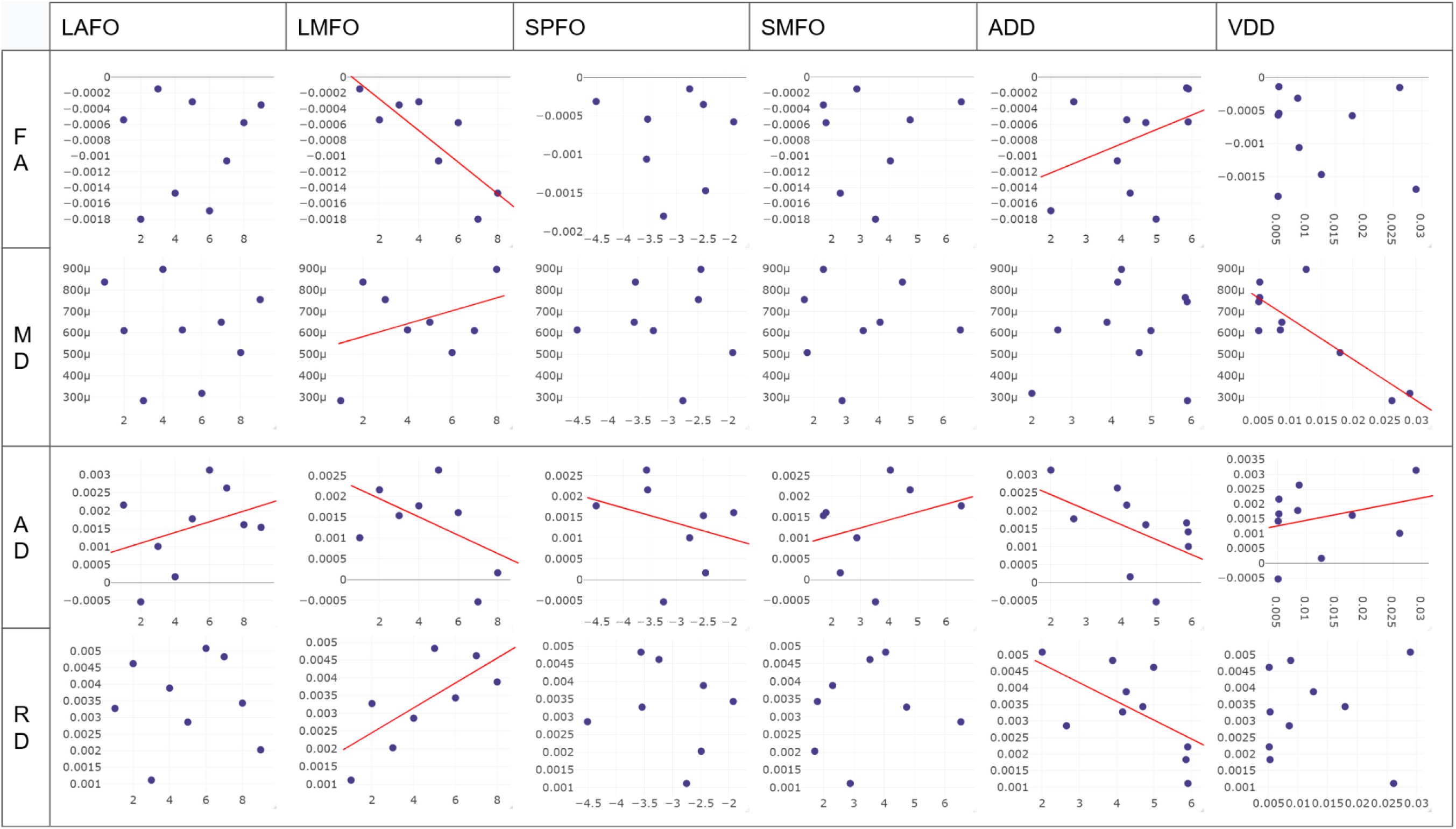
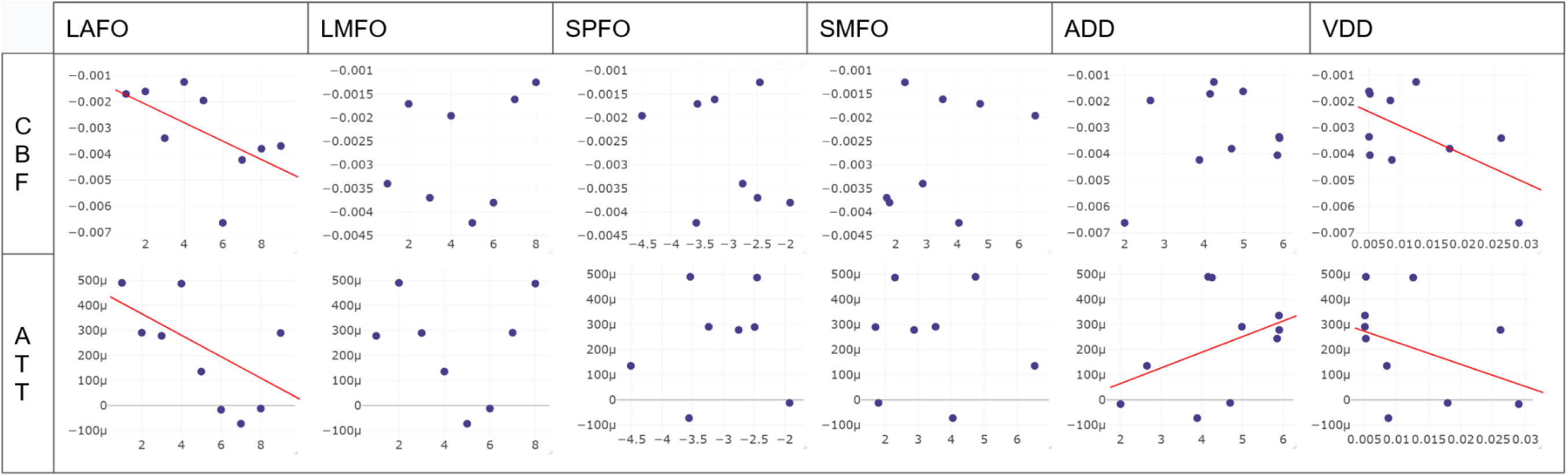
Tractwise age-effects in FA_prec_, MD_prec_, AD_prec_, RD_prec_, CRF_prec_, and ATT_prec_ under each scaled Last-In-First-Out, Gain-Predicts-Loss, and Physiological Determinant model. Dotted fit lines denoting interactions with a p< 0.05, with sex as a covariate.

#### Last-in-first-out (LIFO) models

In the LAFO model, rate-of-decline analyses revealed a significant interaction between tract emergence order and the age effect on AD_perc_; no other significant associations were found. In the LMFO model, tract order demonstrated a negative relationship with the age effect on FA_perc_ and a positive relationship with the age effect on MD_perc_, such that earliest myelinating tracts were associated with slower microstructural deterioration. AD_perc_ and RD_perc_ demonstrated a similar pattern of effects in rate-of-decline interactions to those identified in state-based associations, with myelination order interacting negatively with the age effect on AD_perc_ and positively with the age effect on RD_perc_ (**Figures 4a, d, g**).

#### Gain-predicts-loss models

Tract order in the SMFO model associated positively with the age effect in AD_perc_ in rate-of-decline analysis, i.e. faster myelinating tracts demonstrated the highest rates of diffusivity (suggestive of microstructural decline) with age. Conversely, tract order in the SPFO model was associated negatively with the age effect on AD_perc_, i.e. faster maturing tracts demonstrated the lowest rates of diffusivity increase with age. No significant interactions were identified by FA_perc_, MD_perc_, or RD_perc_ under either model in rates of decline (**Figures 4b, e, h**).

#### Physiological determinant models

In the ADD model, tract order demonstrated a positive relationship with the age effect on FA_perc_, such that higher axon calibre was associated with slower declines in aging. Under the VDD model, a negative association between tract order and age effect on MD_perc_ was found, with higher AxD associated with a higher rate of MD_perc_ increase with age. Axonal calibre was negatively associated with the age effect on both AD_perc_ and RD_perc_ under the ADD model. This pattern is similar to those found in state-based microstructural analyses, where macrovascular density under the VDD model was associated positively with AD_perc_ and was not associated with RD_perc_ (**Figures 4c, f, i**).

### Order of state of perfusion declines

#### Last-in-first-out (LIFO) models

Both the LAFO and LMFO models identified linear relationships between both CBF_perc_ & ATT_perc_. In the LAFO model, order of tract appearance was negatively associated with both CBF_perc_ and ATT_perc_, i.e. the later emerging tracts had both lower CBF_perc_ and and shorter ATT_perc_ later in life. In the LMFO model, CBF_perc_ associated positively with tract order while ATT_perc_ associated negatively, suggesting tracts last to myelinate have higher CBF_perc_ and shorter ATT_perc_ (**Figures 3a**).

#### Gain-predicts-loss models

No significant associations between tract order and mean CBF_perc_ & ATT_perc_ identified by the gain-predicts-loss models (SPFO, SMFO) (**Figures 3b**).

#### Physiological determinant models

Under the ADD model, ATT_perc_ but not CBF_perc_ was found to associate (positively) with axon calibre. Under the VDD mode, higher CBF_perc_ was associated with lower tractwise AxD, while higher ATT_perc_ was associated with lower macrovascular density. No significant associations were found with CBF_perc_ under either physiological model (**Figure 4c**).

### Order of rates of perfusion decline

#### Last-in-first-out models

In the LAFO model alone, significant negative associations were identified between tract order and the strength of age associations with CBF_perc_ & ATT_perc,_ such that later emerging tracts exhibit faster decreases in CBF_perc_ but slower lengthening in ATT_perc_. No interaction between tract-order and age-related CBF_perc_ & ATT_perc_ differences was observed in the LMFO model (**Figure 4a**).

#### Gain-predicts-loss models

No significant interactions between tract order and age effects on CBF_perc_ & ATT_perc_ were identified by the gain-predicts-loss models (SPFO, SMFO) (**Figure 4b**).

#### Physiological determinant models

Under the ADD model, a positive association was identified between AxD and the age effects on ATT_perc_, such that higher age effects were associated with higher axonal calibre. Under the VDD model, a significant positive association between rates of variation in both ATT_perc_ and CBF_perc_, such that higher VasD was associated with faster CBF_perc_ decline but slower ATT_perc_ lengthening (**Figure 4c**).

### Microstructure-perfusions associations

Across the age range, seven of the ten tracts (Fminor, ATR, CAB, CCG, ILF, SLFP, SLFT; not Fmajor, CST or UNC) demonstrated significant associations between WM microstructure and WM perfusion on one or more measures. CBF_perc_ was significantly associated with MD_perc_ in the ILF and SLFT, and with FA_perc_ in the CCG. ATT_perc_ was significantly associated with MD_perc_ in the Fminor, ATR, CCG, SLFP, and SLFT, and with FA_perc_ in the ATR and CAB (**Table 4**).

**Table 4:**
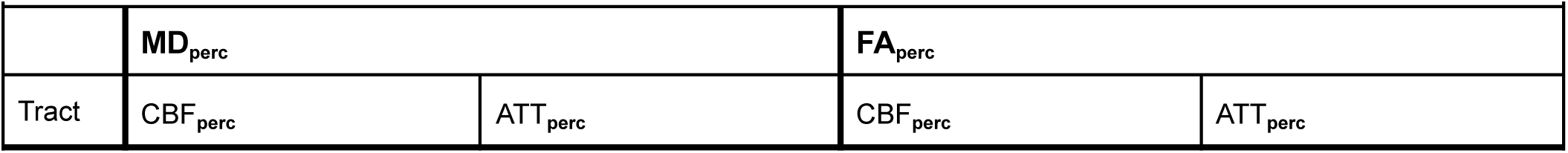
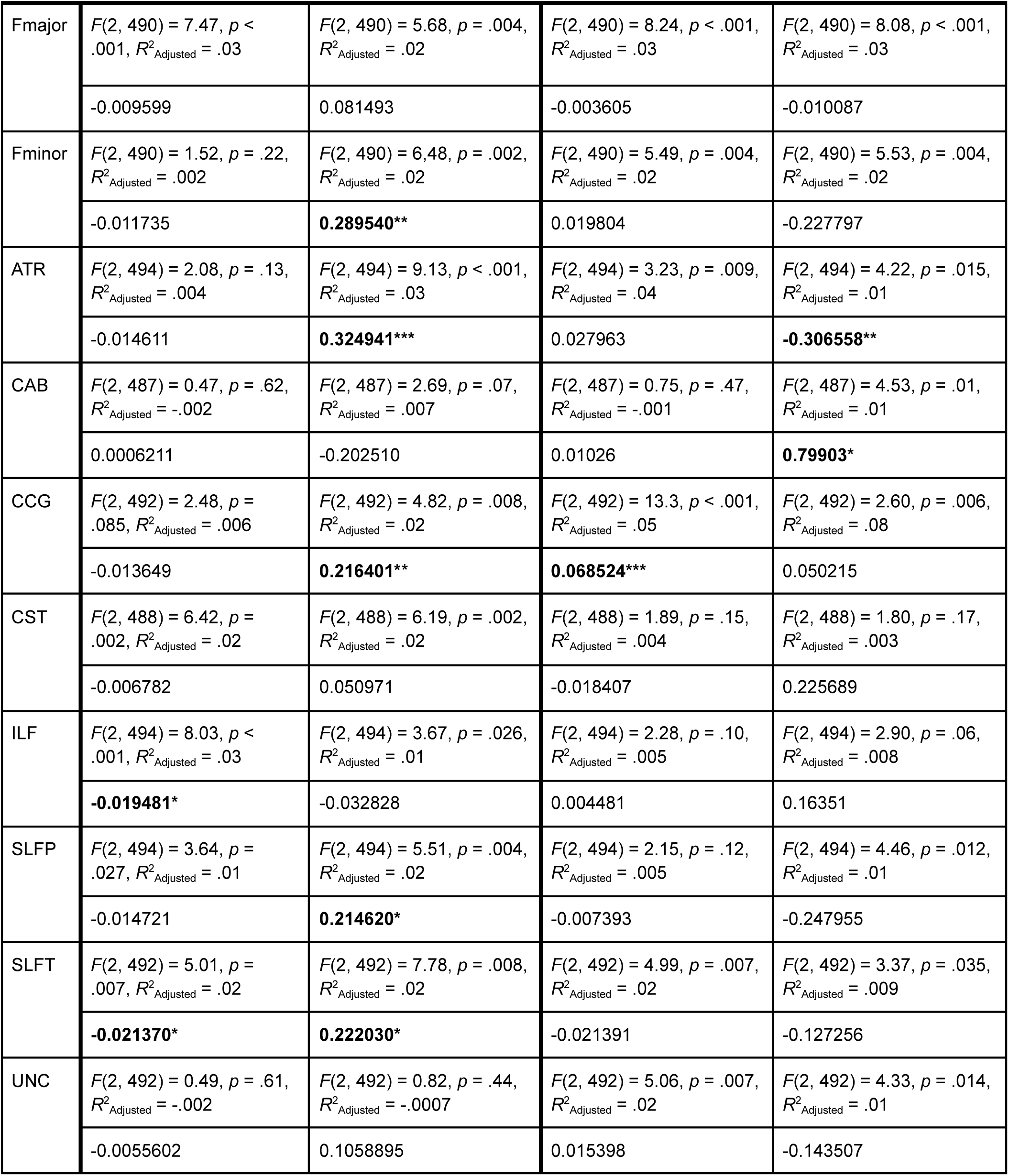
Tractwise MD/FA associations with perfusion (covaried for Sex) All models are multivariate linear regressions based on baseline-normalized measures (x_perc_) with overall F-statistics, *p*-values, and effect sizes (*R^2^_adjusted_*) listed. Slopes are noted below each regression equation and significant associations bolded with asterisks indicating significance. (*:p<.05, **:p<.01, ***:p<.001).

### Baseline associations

#### Inter-subject associations

Significant subject-wise positive associations were identified in baseline values between FA_perc_ and AD_perc_; FA_perc_ and CBF_perc_; MD_perc_ and AD_perc_; and between MD_perc_ and RD_perc_. Negative associations in subject-wise baseline values were found between FA_perc_ and MD_perc_, FA_perc_ and RD_perc_, and between RD_perc_ and CBF_perc_ (**Figure 5a**).

#### Inter-tract associations

Tract-wise positive associations were identified in mean baseline values between MD and AD, MD and RD, and between CBF and AxD (not percent-normalized). Negative associations in tract-wise baseline values were found between FA and RD, MD and CBF, MD and AxD, RD and CBF, and between RD and AxD (**Figure 5b**). Moreover, VasD was not associated with any other baseline metric.

### Effects of sex

Examining the effects of sex on microstructural measures in each of the six tested models identified significant interactions with sex in the association between tract order and microstructural state in four of six models. Female subjects demonstrated stronger associations between tract order and mean FA_perc_ than male subjects under the LAFO, SMFO, and VDD models, and between tract order and both MD_perc_ and RD_perc_ under the ADD model. Conversely, male subjects demonstrated stronger associations between tract order and mean FA_perc_ and RD_perc_ than female subjects under the SMFO and VDD models.

In perfusion state analyses, female subjects showed significantly stronger associations between tract order and CBF_perc_ under the VDD model, and stronger associations with ATT_perc_ than male subjects under the ADD model, while male subjects showed stronger associations between tract order than female subjects in CBF_perc_ under the ADD model, and ATT_perc_ under the LMFO model (**Table 5a**).

**Table 5a:**
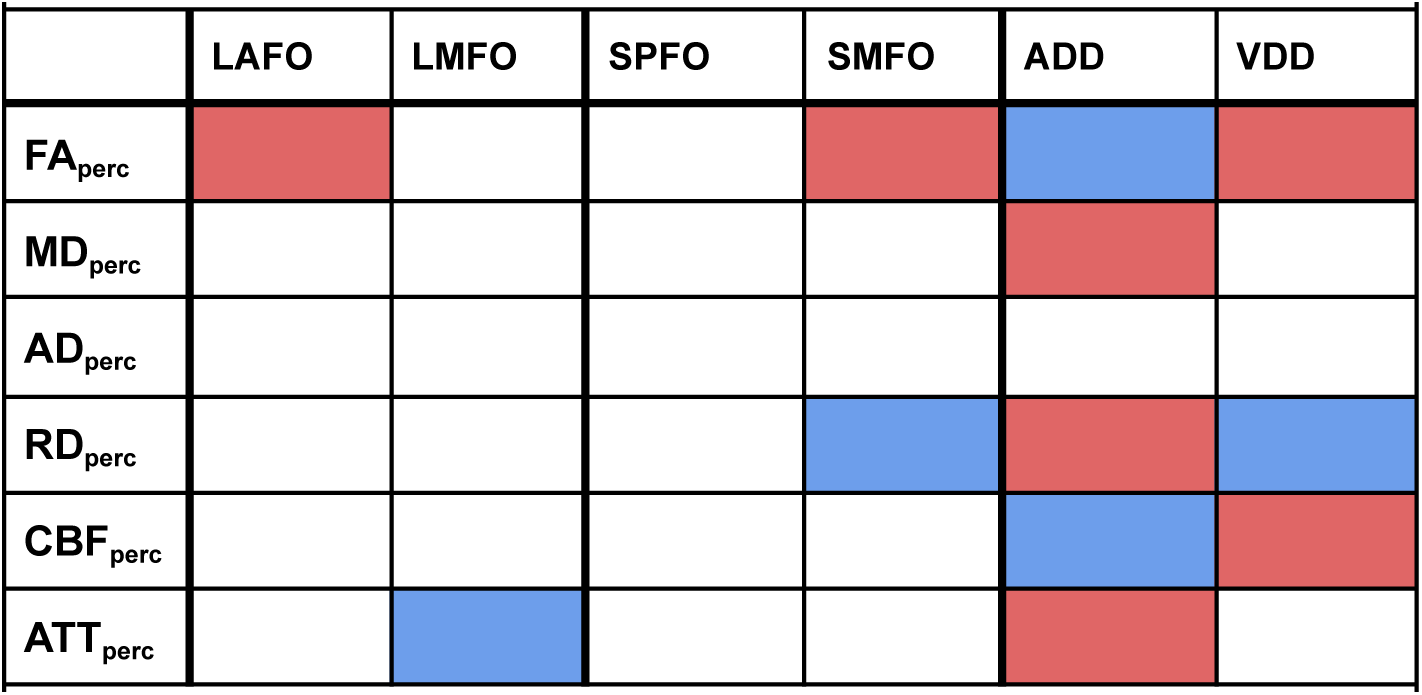
Sex differences (state-based) Color denotes significant (p<0.05) interaction effects of sex on the association between tract order and microstructure perfusion state were identified. Red: higher interaction in females, Blue: higher interaction in males

Two significant interactions were found between subject sex and the effect of tract order on rates of decline in aging. Female subjects showed stronger associations between tractwise macrovascular density (VDD) and CBF_perc_ changes with age than male subjects, while male subjects demonstrated stronger associations between axon calibre (ADD) and rates of AD_perc_ change with age. (**Table 5b**).

**Table 5b:**
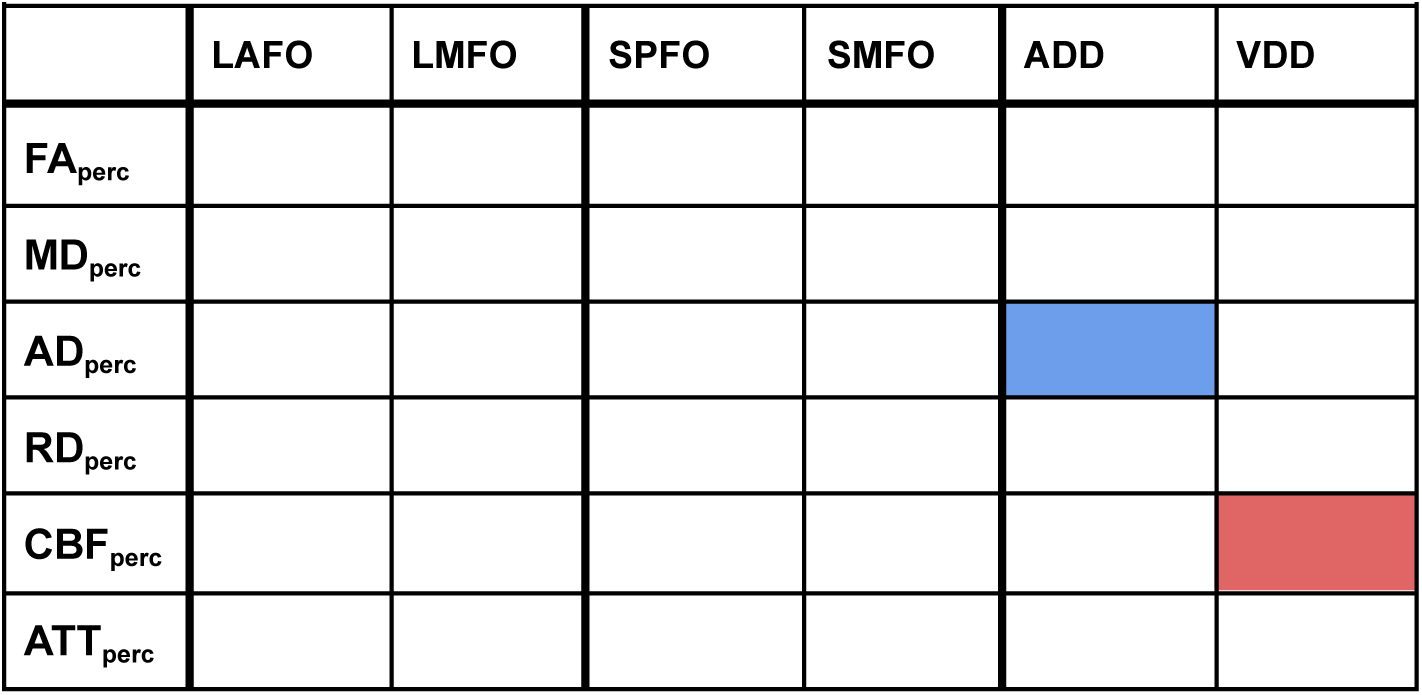
Sex differences (rate-of-decline) Color denotes significant (p<0.05) interaction effects of sex on the interaction of age on the association between tract order and rate of microstructure/perfusion declines were identified. Red: higher interaction in females, Blue: higher interaction in males

## DISCUSSION

The retrogenesis models considered by this study hypothesized that tracts which develop earliest would show the lowest overall declines in adulthood (LIFO), although the definition of “earliest developing” remains ambiguous, necessitating multiple LIFO models. An alternate hypothesis is that tracts which develop most rapidly were expected to exhibit the most rapid declines in aging, i.e. “gain-predicts-loss”. Moreover, we hypothesized that tract-specific physiology could modulate tract degeneration beyond the developmental trajectory. Our results indicate that tracts most vulnerable to age-related degeneration are those which emerge earliest prenatally and display the highest degrees of myelination at birth, with evidence of higher resilience in tracts with higher axon calibre and macrovascular density. Specifically, we find (i) evidence to support the LIFO models, though with a distinction between the LAFO and LMFO models; (ii) negligible support for the gain-predicts-loss models; (iii) substantial support for physiological models of WM aging, with higher axonal calibre (AxD) and higher macrovascular density (VasD) associated with more preserved microstructure. These findings suggest that the prediction of WM aging needs to consider the interactions between physiological processes and LIFO models.

### Order of microstructural decline

The definition of “last-in” has been ambiguous, and the findings of this paper shed some light on the definition of “retrogenesis”, specifically in terms of (1) whether order of prenatal emergence or time to peak lifespan maturation defines “last-in”; (2) whether “last-in-first-out” or “gain-predicts-loss” better defines the retrogenesis trajectory.

Interestingly, the LAFO model identified a trend whereby later-emerging tracts have higher FA and AD in the state-based analysis, indicative of higher integrity. Given that the order of tracts presented by the LAFO model follows a generally anterior to posterior pattern of tract emergence, this would imply that posterior WM maintains higher integrity (Hoagey et al., 2019; Slater et al., 2019). Conversely, as the LMFO model presents tracts in the opposite order (i.e. WM tracts mature in the opposite order of their initial emergence), the tracts myelinated earlier maintain higher integrity. Between the two models, the LMFO model is a better predictor of WM integrity than the LAFO model. As these ordinal models reflect different components of prenatal development, with existing evidence supporting opposing orders of development between first appearance of tract structures and their degree of myelination at birth (Grotheer et al., 2022; M. Ouyang et al., 2019), this finding illustrates the complexity in the process of “genesis” and hence the definition of retrogenesis. Factors other than structural or microstructural integrity, including angiogenesis and glial maturation (Arai et al., 2018; Hanslik et al., 2021; Hodges et al., 2018; Soreq et al., 2017), are likely integral to these definitions. Conversely, the lack of significant associations between SPFO and SMFO tract orders and FA_perc_ or MD_perc_ states suggests a lack of support for the theory that fibres that develop slower degenerate faster (gain-predicts-loss) (Bartzokis, 2004).

On the other hand, the associations identified by the ADD and VDD models indicate support for thinner, less myelinated and less vascularized axons being more vulnerable to degeneration in terms of both FA_perc_ and MD_perc_. These are also tracts with less myelin, as reflected by their higher R1 values (used in the LMFO model) (Bartzokis, 2004).

### Order of perfusion declines

Continuing the developmental patterns observed in microstructure, a greater rate of CBF_perc_ decline was associated with later tract emergence. As declines in CBF are assumed to indicate a greater degree of age-related microvascular impairment (Bernbaum et al., 2015; Chen et al., 2013), this finding would support retrogenesis when defining “first-in” in terms of the order of tract emergence. Increasing ATT would similarly be assumed to indicate age-related microvascular declines, and ATT_perc_ demonstrates less change with age in the later emerging tracts. That is, the later a tract emerges in prenatal development the more rapidly CBF_perc_ declines but the less rapidly the ATT_perc_ lengthens in aging. CBF and ATT are thus shown to not mirror each other in aging.

This dichotomous association between CBF and ATT variations as a function of tract development is very interesting and has been previously identified in aging haemodynamics (Yetim et al., 2023). This pattern may suggest differences in regional compensatory mechanisms in microvascular decline across early- and late-developing regions. That is, it is possible that early-developing regions enjoy relatively maintained overall CBF though it is increasingly delivered with an collateral routes resulting in increasing ATT (Maguida & Shuaib, 2023), whereas later-developing regions may not be able to function in this regime, either due to greater demand for blood flow or a lack of effective collaterals.

The vascular endothelial growth factor, which drives angiogenesis and the formation of collateral flow routes, is highest in early brain development (Maharaj & D’Amore, 2007; Rosenstein et al., 2010), perhaps resulting in more robust vascular supply for earlier-forming tissue regions. The possibility also remains that early WM microvascular development in childhood and adolescence may vary considerably with the order of development, with the vascular routes to early-developing regions expanding more slowly than later developing regions. For instance, frontal regions, despite the rapid CBF increase in development (M. Ouyang et al., 2019), lack the leptomeningeal collateral circulation routes that are possessed by the more posterior brain regions (Hung et al., 2022).

This pattern was also associated with axon calibre under the ADD model. Ex-vivo tissue development studies suggest that axon development follows vascular development, with higher density axon development in proximity to larger vascular sources (Pawar et al., 2015), while axon calibre may instead depend on the activity level of associated cortical tissue (Perge et al., 2012). As our data demonstrates, earlier-developing tracts are associated with larger axonal calibre, which is in turn associated with more rapid ATT lengthening in aging as well as lower macrovascular density, it may be reasonable to surmise that larger axons enjoy collateral flow in aging and are thus spared from more severe WM degeneration. With that in mind, a lengthening ATT may not always indicate degeneration. This is an important finding that could alter the interpretation of ATT in the study of brain aging. However, it remains unclear if the larger axons survive longer because of the presence of collateral flow or a lower metabolic demand, highlighting the need to further our understanding of changes in metabolic demand across the lifespan and their influences on neurovascular development and deterioration.

In contrast to the LAFO model, tract ranking based on the LMFO model was negligibly related to CBF or ATT variations in aging. However, as noted earlier, previous literature has demonstrated that the developmental orders in tract emergence and myelination in-utero contrast one another (Grotheer et al., 2022; Y. Ouyang et al., 2021). Additional examination of early prenatal development, along with the previously suggested opposing orders of prenatal tract emergence order and myelination, is required to clarify how the earliest stages of prenatal development interact. Although retrogenesis in terms of perfusion decline may favour the last tract to emerge as “last in”, we cannot conclude this definitively as we do not have developmental perfusion trajectories. Lastly, perfusion variations were not determined by “gains-predict-loss” as defined by WM development.

Perfusion results deviate from patterns identified in microstructural declines when examining other models, and across measures, state-based analyses largely corroborated the patterns found in rate-based mean microstructural states. An exception was identified under the LMFO model, where neither CBF_perc_ or ATT_perc_ rates were associated with myelination levels at birth, but tract-wise differences were found in perfusion states. The lack of strong ties between peak-myelination order and perfusion variables across tracts implies that CBF decline is not a mirror image of CBF growth in development, because we assume that earlier peak myelination reflects more rapid myelination and thus may require more rapid angiogenesis (Tsai et al., 2016; Yuen et al., 2014).

### Microstructure-perfusion associations

#### Role of developmental trajectory

With the majority of tracts confirming the presence of associations between microstructure and perfusion, it can be assumed that both do, to an extent, reflect a common mechanism of age-related declines. While the results from direct microstructure-perfusion associations can be taken as support for common mechanisms of declines across microstructural and perfusion measures, these associations were not universal. Associations varied by metric between tracts and three of the tracts examined (Fmajor, CST, and UNC) did not demonstrate microstructure-perfusion associations on any metric. Moreover, it is notable that microstructural and perfusion declines do not follow the same order across tracts, and are not equally predictable from the same models of retrogenesis. There was consistent association between the rate of perfusion decline and tract emergence order under the LAFO model but not the LMFO model, whereas microstructural decline was more consistently associated with the LMFO than the LAFO model. That is, earliest prenatal emergence may be more strongly associated with vascular development, while prenatal myelination associates most consistently with microstructural integrity across analyses, suggesting the presence of diverse compensatory mechanisms that support tract-wise resilience to age.

#### Physiological associations at baseline

Across participants, a negative association was found between RD_perc_ and CBF_perc_ within the baseline group, suggesting individuals with lower CBF would also manifest higher RD, even without the aging factor. This is consistent with our understanding of the importance of CBF in maintaining WM microstructural integrity (Chen et al., 2013; Robinson et al., 2023). However, despite negative trends in the baseline association between CBF and both MD_perc_ and RD_perc_, these associations were not significant across tracts. ATT was not involved in this association, suggesting that variations in ATT may not be as critical for predicting WM integrity. Across tracts, the only significant relationship between the baseline physiological variables and our microstructural measures of interest was a negative association between axon calibre and RD_perc_, suggesting larger axons being associated with lower RD_perc_, potentially due to both higher integrity and tighter myelin packing (Osso & Hughes, 2024).

The lack of relationship between WM CBF and macrovascular density may indicate that macrovascular density is not a strong predictor of regional microvascular physiology. As such, incorporating other vascular variables into these predictive models, ones such as ATT and the number of regional vascular sources, may be necessary.

#### Tying things together

Across analyses, our results show that both early developmental staging and physiological differences are linked to patterns of WM decline in aging. Evidence of differences in axon calibre and myelin content between tracts and sexes across childhood and adolescence (Genc et al., 2018) suggest that physiological and developmental measures are interrelated, with axon calibre varying by tract across age. As such, developmentally ordered models may inadvertently capture tractwise differences in physiology. In our work, WM decline rate was significantly associated with both axonal calibre and vascular density, but these latters were not significantly associated with developmental trajectory across tracts or across participants. Moreover, the definition of the WM development trajectory itself may be up for debate as it relates to the idea of “retrogenesis”, as the trajectory as defined by tract emergence and myelination appear to contradict each other in their relationships with WM decline.

Furthermore, our results suggest that our understanding of the relationships between microstructural and microvascular declines should be founded on tract-wise differences in physiology. For example, if a physiological difference such as higher axon calibre were to result in a region being less metabolically demanding and thus more resilient to perfusion declines, it could explain the lack of strong associations between microvascular health and microstructural declines that are detectable in less resilient regions. How developmental order and myelination, physiological differences in WM regions, and vascular supply interact with regards to the resilience of WM tissue and underlying metabolic demand likely presents a more comprehensive and accurate model of WM declines in aging than each individual model alone. Clarifying these interactions will require a more complete understanding of the developmental and hormonal drivers of physiological development by tract.

### Sex-related differences

Extensive sex differences in rates and states of microstructural and perfusion declines across age were observed under the ADD and VDD models, though no single pattern across all models was identified. Under the ADD model, AD_perc_ rate of change was more significantly (and negatively) associated with axonal calibre in males, whereas under the VDD model, CBF_perc_ rate of change is more strongly (and negatively) associated with VascD in females. These observations may suggest that either males demonstrate a greater range of axon calibres between tracts, or otherwise that males display greater differences in regional susceptibility to WM declines. Likewise, females may demonstrate a greater range of CBF values or VascD across tracts. Sex differences in rate of WM decline were not observed in relation to the developmental models, but were observed for the state of WM integrity under the LAFO and SMFO models. We note that some studies have found higher myelin density in males during WM development (Corrigan et al., 2021), but the findings have in general not been consistent (Cercignani et al., 2017). In general, the majority of the sex differences are found under the physiological models, which illustrates the need for a better understanding of the causal relationships between macro- and microvascular health as they relate to microstructural integrity in age. Moreover, the ADD and VDD models demonstrate the opposite sex differences, which is consistent with the mild but negative slope between baseline AxD and VascD (**Fig. 6b**).

**Figure 6:**
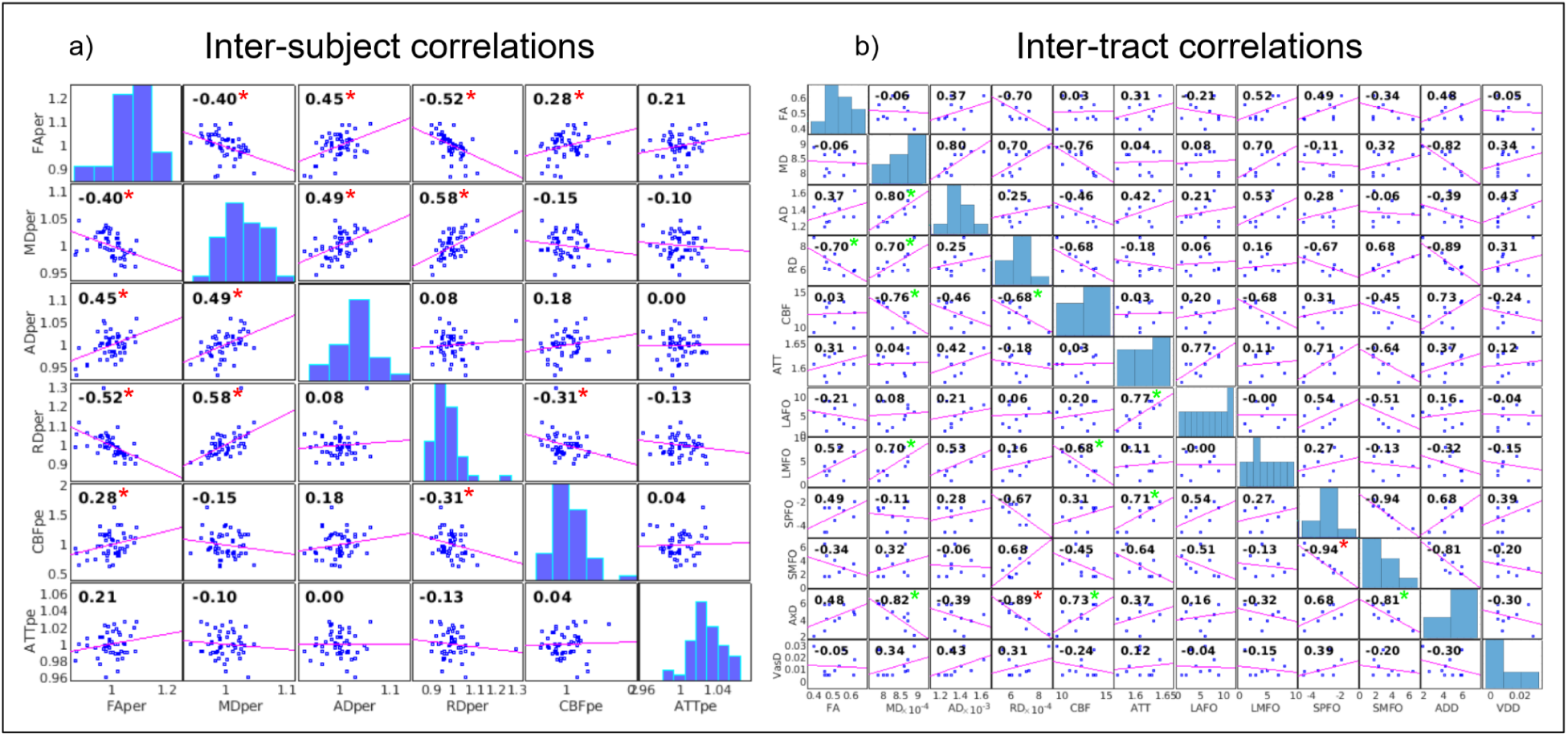
Baseline relationships between tai inter-subject microstructural (FA _prec_, MD_prec_, AD_prec_, RD_prec_) and perfusion (CBF_prec_, ATT_prec_) measures and (b) inter-tract microstructural (FA, MD, AD, RD), perfusion (CBF, ATT), developmental orders (LAFO, LMFO), maturation rates (SPFO, SMFO), and physiological (AxD, VascD) measures. Correlations withp<0.05 ate denoted with asterisks in red.

While existing research on the topic remains limited, previous evidence in animal studies of corpus callosum development suggest that both of these types of sex-related differences may be present (Aboitiz et al., 1996; Pesaresi et al., 2015; Shatarat et al., 2020). Additionally, testosterone has been found to regulate axon radius, leading to higher mean axon calibre in the splenium of male than female rats (Pesaresi et al., 2015). Thus, based on our previous finding that females exhibit greater rates of WM decline (Robinson et al., 2024), a follow-up question related to the findings under the ADD model is whether the lower axonal calibres in females predisposes females to WM decline. While sex hormones are known to be involved in the prevalence of and susceptibility to many neurovascular conditions, as well as having suggested roles in vascular autoregulation (Dion-Albert et al., 2022), sex differences in vascular density in the white matter remain largely understudied. Recent exploratory evidence has identified sex differences in arterial density across the brain, with female subjects showing lower values in the periventricular WM that may result in greater vulnerability to WM hypoperfusion in female subjects (Barbeau-Meunier et al., 2022).

### Limitations

The ordered models applied in this study are drawn from a number of prior sources examining developmental differences in the WM. As such, this examination has been limited to those studies examining similar tracts to allow for cross-model comparisons using similar reconstructed tracts. These tracts were chosen to minimize tract overlap/partial-voluming between segmentations, however, additional comparisons may be possible in more recent segmentations (Maffei et al., 2021). For instance, future work can benefit from further segmenting tracts in terms of anterior versus posterior segments.

Secondly, as the LAFO and LMFO models applied by this study are ordinal in nature due to limited availability of quantitative data on developmental differences, they remain limited in their ability to accurately represent fine differences in regional development that would be possible with quantitative models.

Thirdly, this study chose to address microstructural measures such as MD and FA commonly applied in existing research. This introduces limitations that may not be present with more advanced diffusion modeling.

Fourthly, the HCP-A sample examined in this study represents a cognitively healthy sample extending well into older age. While Brookheimer et al. have addressed the necessary presence of limited degrees of cognitive and vascular declines in a sample of this age, by selecting for cognitively sound older subjects, the sample may represent an atypically well-aging population in addition to being more highly educated than average for the US population (Bookheimer et al., 2019).

Moreover, while our between-tract analysis demonstrates meaningful patterns within both this study and previous research (Robinson et al., 2023), within-tract comparisons represent an additional axis of comparisons that has received relatively little attention. Given the considerable differences in rates and patterns of development across tracts in all stages of WM growth and maturation, comparing regions within tracts along a developmental gradient may further clarify our understanding of retrogenesis as a process.

Finally, the cross-sectional nature of the HCP-A dataset used here could be complemented by future examinations using longitudinal data.

## CONCLUSIONS

In addition to clarifying the stages of WM development that are predictive of later aging declines, there remain gaps in our understanding of how developmental and physiological factors anticipate patterns of WM aging. This study demonstrated support for the retrogenesis hypothesis in the form of developmental differences as a predictor for tract-wise WM microstructural and perfusion declines in aging. Also, the order of prenatal tract emergence was found to have an inverse relationship with the order of aging declines in WM, opposite that asserted by late-in-first-out predictions. Additionally, both axon calibre and macrovascular density were found to be significant physiological predictors for tract-wise aging differences, with regions of higher mean axon calibre and lower macrovascular density exhibiting higher resilience to age-related degeneration. Importantly, these physiological influences on WM decline are much more sex dependent than developmental influences, which merits further study. To our knowledge, this study is the first to demonstrate the major role of physiology, in addition to developmental order, in explaining the patterns of age-related WM decline.

## Acknowledgments

We are grateful for the financial support from the Canadian Institutes of Health Research (CIHR) (#PTJ169688) and the Canada Research Chairs (CRC) program (JJC). Research reported in this publication was supported by the National Institute On Aging of the National Institutes of Health under Award Number U01AG052564 and by funds provided by the McDonnell Center for Systems Neuroscience at Washington University in St. Louis. The HCP-Aging 2.0 Release data used in this report came from DOI: 10.15154/1520707.

